# VIPER: A General Model for Prediction of Enzyme Substrates

**DOI:** 10.1101/2024.06.21.599972

**Authors:** Maxwell J. Campbell

## Abstract

Enzymes, nature’s catalysts, possess remarkable properties such as high stereo-, regio-, and chemo-specificity. These properties allow enzymes to greatly simplify complex synthetic processes, resulting in improved yields and reduced manufacturing costs compared to traditional chemical methods. However, the lack of experimental characterization of enzyme substrates, with only a few thousand out of tens of millions of known enzymes in Uniprot having annotated substrates, severely limits the ability of chemists to repurpose enzymes for industrial applications. Previous machine learning models aimed at predicting enzyme substrates have been hampered by poor generalization to new substrates. Here, we introduce VIPER (Virtual Interaction Predictor for Enzyme Reactivity), a model that achieves an average 30% improvement over the previous state-of-the-art model (ProSmith) in reaction prediction for unseen substrates. Furthermore, we reveal flaws in previous enzyme-substrate reaction datasets, and introduce a novel high-quality enzyme-substrate reaction dataset to alleviate these issues.

## 1 Introduction

Enzymes, nature’s catalysts, possess unique catalytic properties, including high stereospecificity, regiospecificity, and chemospecificity (1). These properties allow for the simplification of complex synthetic processes including ones used in the synthesis of several FDA-approved drugs (1,2). One recent example is the biosynthesis of QS-21 (3), a potent immunostimulatory agent used in several FDA-approved vaccines (4). While the chemical synthesis of the Api and Xyl components of QS-21 requires 76 steps (5–7), the biosynthesis only requires 20 steps (3), significantly improving the yield and potentially significantly decreasing the cost of manufacturing QS-21. Additionally, enzymes have great potential for use as eco-friendly catalysts as they operate under mild reaction conditions, such as ambient temperature and pressure making them an eco-friendly choice for industrial processes (8).

However, very few enzyme substrates have been experimentally characterized, with only a few thousand out of tens of millions of enzymes in Uniprot (9) having experimentally confirmed substrate annotations, making repurposing previously discovered enzymes for use in industrial reactions difficult, as the enzyme substrates must be manually characterized in high-throughput screens, a time-consuming and expensive endeavor. This limitation hampers the ability of chemists to select an enzyme off the shelf as they would a chemical reagent for a traditional reaction, making it difficult to fully realize the potential of enzymes in industry.

Previous works have attempted to address this issue by introducing broad machine learning models for the prediction of enzyme substrates. Three models for this task have been introduced previously: ESP (10), ProSmith (11), and Ridge Regression (12). ESP is based on XGBoost (13) and uses data scraped from GO annotations (14). Fine-tuned ESM-1b (15) embeddings are utilized to embed the protein and Graph Neural Network (GNN) embeddings were utilized to embed the molecule. While ESP is capable of learning to predict enzyme-substrate reactions for the substrates included in its training set, as we will discuss later it performs poorly for distant substrates achieving an MCC of 0.320 on distant unseen substrates. ProSmith also uses data scraped from GO annotations, it is a hierarchical combination of a pre-trained transformer model (16) that takes in ChemBERTa2 (17) and ESM-1b embeddings, then feeds the output into an ensemble of XGBoost heads, along with the original ChemBERTa2 and ESM-1b embeddings. ProSmith was pretrained on a kinase inhibitor binding affinity prediction task, using the Davis dataset (18). ProSmith achieves an MCC of 0.314 on our unseen substrate benchmark. Finally, the Ridge Regression model uses data from high-throughput screens with ESM-1b and Extended-connectivity fingerprint (ECFP) embeddings. The Ridge Regression model achieves an MCC score of 0.334.

Other work has introduced models for the prediction of enzyme substrates within singular families. Several models have been developed for this purpose; we will highlight two here. Mou et al. developed a decision-tree-based model that predicts the substrates of bacterial nitrilases using substrate-protein 3D model-based descriptors trained on 12 nitrilases and 20 nitriles (19). The other model proposed by Yang et al. also uses a decision tree approach with manual chemical and enzyme features trained on 54 glycosyltransferases and 94 substrates (20). Unfortunately, like the previously discussed models, these models only achieve good performance within the enzyme family they are trained on and do not learn general enzymatic properties or reaction schemes (10).

There have been some inroads made into the prediction of Enzyme Commission (EC) numbers for enzymes. Models such as CLEAN (21) and DEEPre (22) have had great success in grouping unannotated enzymes into families. Unfortunately, they are not useful for profiling the activity of enzymes toward specific substrates beyond knowing what family the enzyme lies in.

While previous models have taken significant steps towards achieving the ideal of enzyme-substrate prediction, as we will show later they either do not generalize well to new substrates (10–12), or are too broad to be of practical use when the family an enzyme lies in is already known (21,22). A model’s practical use is dependent on its ability to be interpreted from a physicochemical standpoint, rather than relying on hidden patterns in the training data or rote memorization (23); therefore, we validated that our model learns generalizable enzyme-substrate reaction mechanics by evaluating it’s out-of-distribution generalization capabilities, and by performing an ablation study to determine the effect of cross-attention, a compound-protein interaction (CPI) mechanism, on VIPER’s performance.

In this work, we introduce VIPER (Virtual Interaction Predictor for Enzyme Reactivity), a machine learning model that outperforms previous models in predicting enzyme-substrate reactions on unseen substrates and exhibits superior out-of-distribution generalization. In addition to evaluating VIPER on unseen substrate performance we conduct a thorough investigation of VIPER’s out-of-distribution generalization capabilities and compare its performance to previously introduced enzyme-substrate prediction models. Additionally we introduce new techniques for evaluating the quality of enzyme-substrate reaction datasets, and show that previous datasets are flawed. Furthermore, we introduce an enhanced dataset for enzyme-substrate interactions, building upon the work of Goldman et al. (12) by incorporating additional data cleaning and standardization techniques to ensure high-quality data.

## 2 Results

### 2.1 Model architecture and training strategy

Enzyme-substrate prediction is a challenging task due in part to the limited amount of available data. To address this issue, VIPER employs a transfer learning approach by utilizing pre-trained transformer models to generate embeddings for both the input molecule and protein. This allows the model to leverage knowledge about proteins and molecules acquired during pre-training and apply it to enzyme-substrate prediction. For protein embedding, VIPER utilizes Ankh base (24), a 450 million parameter protein language model. Ankh base was pre-trained on the UniRef50 (9) database, which consists of 45 million protein sequences, enabling it to capture a wide range of protein characteristics and features. For molecule embedding, VIPER employs Molformer (25), a 44 million parameter chemical transformer model pre-trained on a dataset consisting of 10% of ZINC (26) and 10% of Pubchem (27), amounting to roughly 110 million molecules. This extensive pre-training allows Molformer to learn a rich representation of molecular structures and properties.

Our model then takes the Ankh and Molformer embeddings as input, which are subsequently processed by separate 1D Convolutional Neural Networks (CNNs). The CNNs utilize the structured input to extract meaningful features from the input embeddings. The CNN outputs are then fed into a cross-attention layer (16); the cross-attention mechanism helps VIPER learn interactions between the molecule and protein. After cross-attention is applied, the resulting protein and molecule embeddings are concatenated together. This combined representation is then fed into a bidirectional Long Short-Term Memory (LSTM) layer. The LSTMs ability to capture long-range dependencies allows the model to learn interactions between distant protein and molecule features. The output of the LSTM is then fed into an Multi-Layer Perceptron (MLP). The final output is a prediction score between 0 and 1, indicating the likelihood of interaction.

To optimize the model, VIPER combines the AdamW optimizer (28) with Sharpness Aware Minimization (SAM) (29). SAM is used in conjunction with AdamW to encourage the model to converge to flatter minima in the loss landscape. Flatter minima are associated with better generalization performance, as they are less sensitive to small perturbations in the input data (29,30). This is particularly important for enzyme-substrate prediction as the data is inherently noisy, as some degree of experimental variance is always expected. To handle the imbalanced nature of the dataset, VIPER uses a weighted binary cross-entropy loss function. The weights are set to a 0.3:1 ratio, giving more importance to the minority class (Positive reactions). This helps prevent the model from being biased towards the majority class and ensures that it learns to correctly classify both positive and negative samples.

### 2.2 A curated dataset from high-throughput screens

We compared enzyme-substrate predictions models on a dataset consisting of nine high-throughput enzyme substrate screens extracted from the literature. This dataset consisted of a relatively diverse set of enzyme families, including phosphatases (31), esterases (32), glycosyltransferases (20), beta-keto acid cleavage enzymes (33), thiolases (34), halogenases (35), aminotransferases (36), and nitrilases (37). Using this experimental data provided several key advantages over relying on datasets derived from GO annotations (14) or the BRENDA database (38).

One of the most significant advantages of our dataset lies in its ability to avoid misannotations, which are pervasive in human-curated annotation databases. Numerous studies have documented the prevalence of misannotations across various databases, including KEGG, BRENDA, UniProtKB, and GenBank NR (39,40). In some cases, misannotation rates have been reported to exceed 75% across different EC numbers and enzyme families (39,40). Previous studies focused on enzyme-substrate prediction have sought to address these issues by implementing heuristics and rule-based systems to clean the data (41). However, the effectiveness of such methods is limited when dealing with data characterized by such high error rates. A particularly striking example is presented by Rembeza et al. (40), where they analyzed 122 representative sequences of the EC number 1.1.3.15 in the BRENDA database and discovered that at least 78% of the data in this family was incorrectly annotated. By directly utilizing data obtained from high-throughput enzyme-substrate screens, our approach effectively circumvents these challenges.

Our dataset offers another key advantage in that it includes negative enzyme-substrate pair reactions, owing to its direct acquisition from high-throughput enzyme-substrate screens. Such information is frequently absent in annotation databases, which has resulted in previous work (10,11) artificially creating negative samples resulting in reduced data quality. For instance Kroll et al. (10) generated negative samples by selecting molecules from their dataset that are similar to a known substrate for the given enzyme. These molecule-enzyme combinations are then added as negative records. Kroll et al. argue that the enzymatic and molecular diversity of their dataset and the combinatorial complexity that arises from it, makes it unlikely that one of the selected molecules will catalyze a reaction with the given enzyme. This argument is flawed as the dataset does not exhibit the assumed diversity. When EC numbers are truncated to their first three digits (e.g., EC 1.1.1.x → EC 1.1.1) to group enzymes by reaction type while avoiding potential misclassifications at more specific levels, 71% of records are concentrated in just 25 out of 187 EC classes (Supplementary Figure 2). Similarly the molecular space is not randomly sampled but instead clustered around the known substrates of these enzymes. This makes the probability of accidentally selecting a true substrate as a negative example much higher than they estimate. This issue is particularly evident in the results of Kroll et al. where they show significantly decreased performance for molecules not present in their training set, even when testing on enzymes with high sequence similarity to their training data (10). Moreover, high-throughput screens generate data with substantially lower sparsity (i.e., fewer missing enzyme-substrate pairs) and decrease experimental variance by assessing enzymes against a wide range of substrates under controlled conditions. In contrast, annotation datasets often suffer from high sparsity and high experimental variance.

To facilitate quantitative comparison between our dataset and that of Kroll et al. (10), we developed two novel metrics for assessing enzyme-substrate dataset quality. The first metric quantifies the prevalence of instances where a common cofactor (e.g., NAD, ATP) is recorded as the primary substrate rather than the enzyme’s actual substrate of interest. Given that current enzyme-substrate prediction models are constrained to single-substrate predictions, the selection of the mechanistically relevant substrate over ubiquitous cofactors is crucial for maintaining dataset fidelity. The second metric evaluates reaction feasibility by leveraging the RetroRules database (42). For each enzyme-substrate pair, we identify the corresponding EC number and attempt to apply all associated reaction rules from RetroRules to the substrate molecule. Reactions are classified as infeasible when no valid transformation rules exist for the given molecule-EC number pair, providing a systematic approach for identifying potentially erroneous annotations.

As evident from the dataset comparison in Figure 2, the Kroll et al. dataset (10) exhibits a 11.1% rate of lone cofactors and a 19.9% rate of impossible reactions, leading to a cumulative error rate of 31.0% for positive interactions. In contrast, the VIPER dataset has a 0.0% rate of lone cofactors and a 9.3% rate of impossible reactions, resulting in a total error rate of 9.3% for positive interactions.

**Figure 1.**
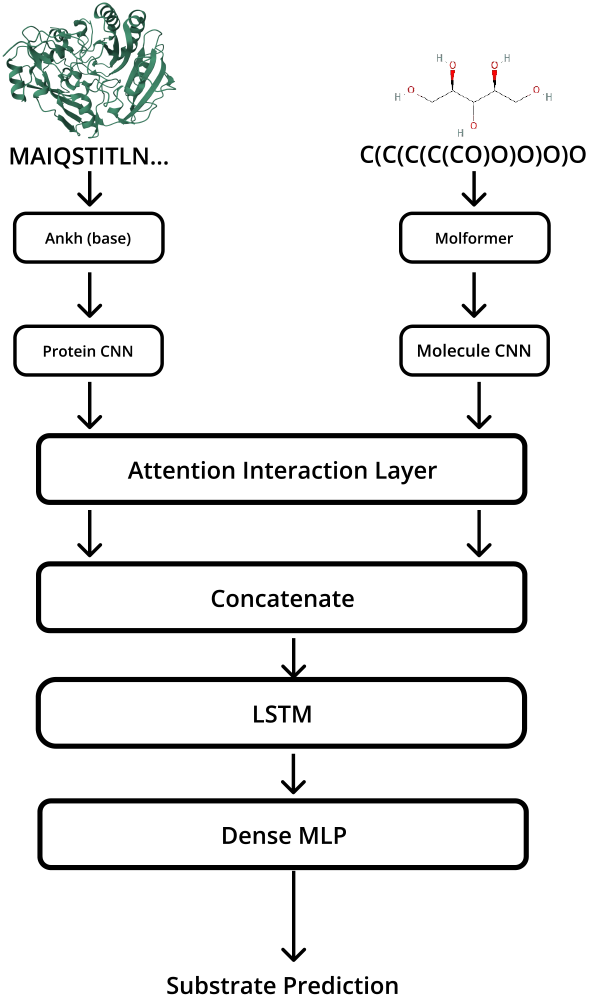
The protein’s amino acid sequence is embedded using the pre-trained protein language model, Ankh (24), and the molecule’s SMILES string is embedded using Molformer (25). These embeddings are then passed through two separate CNNs, after which the embeddings are fed into a cross-attention layer (16); after this, the tensors are concatenated and fed into an LSTM the resulting tensor is then provided to an MLP to produce a prediction.

**Figure 2.**
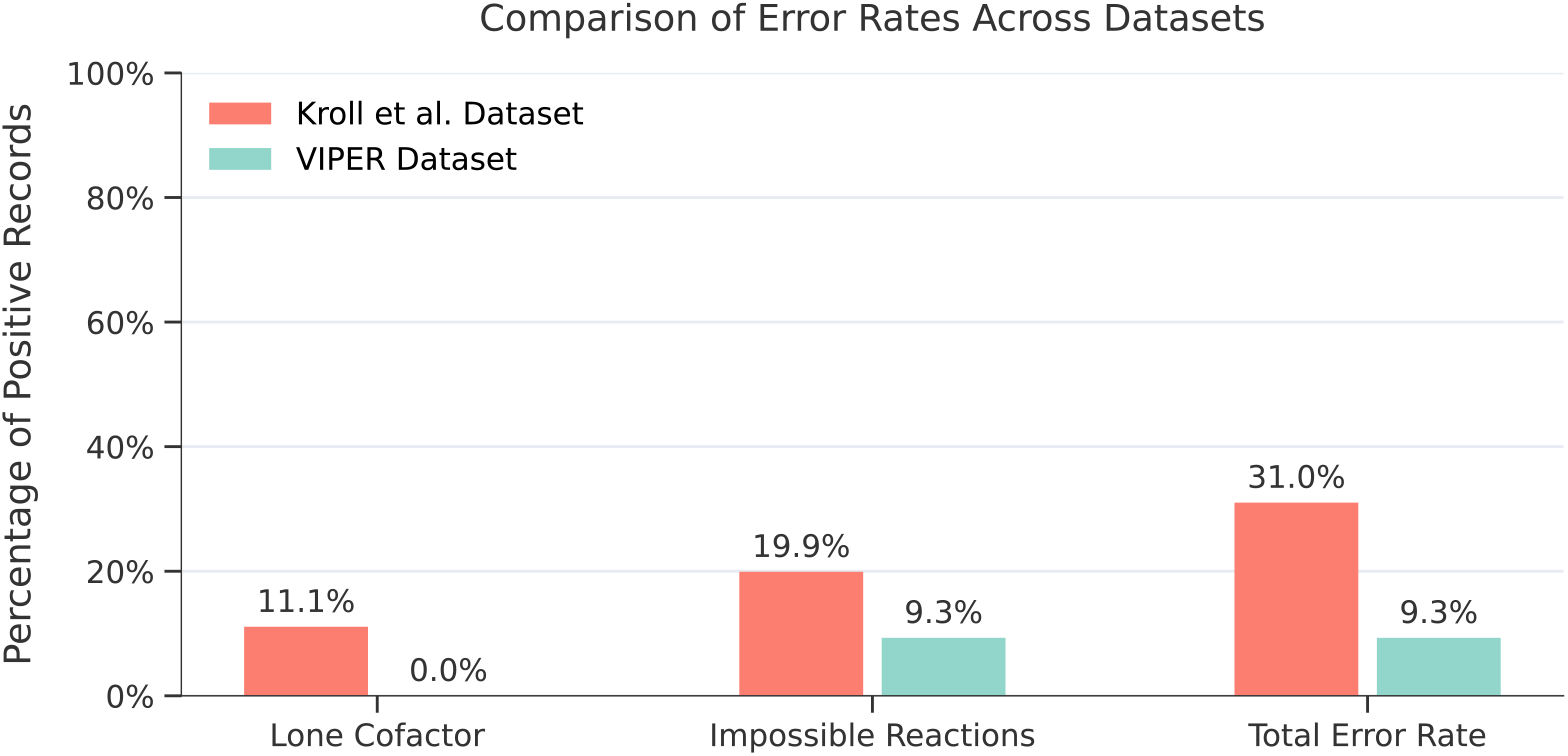
Lone cofactor and impossible reaction error rates compared across VIPER dataset and Kroll et al. dataset (10). The comparison shows three metrics: lone cofactor rates where the substrate is a common cofactor, impossible reaction rates where no valid reaction rule exists for the substrate-EC number pair according to RetroRules (42), and the combined total error rate. Values are expressed as percentages of positive records in each dataset.

### 2.3 Improved generalization to unseen substrates with VIPER

To evaluate VIPER’s performance on unseen distant molecules, we employed a sphere exclusion split on molecular extended connectivity fingerprints (ECFPs) (43) to create a training set, validation set, and testing set with dissimilar molecules. The sphere exclusion method ensures that the test set molecules are sufficiently distant from the training set molecules by enforcing a minimum Euclidean distance between their ECFP representations. For our experiments, we set the Euclidean distance threshold to 0.25, the default configuration in astartes (44). We chose to use euclidean distance to split the dataset as it is a simple extrapolative splitting method that can easily be applied to arbitrary vectorized molecules.

For VIPER, we used the validation set to select the best performing model out of ten training epochs, while for ProSmith, it was used to determine the optimal weighting between three gradient boosting tree models. For ESP, the validation set was used to perform hyperparameter optimization as described by Kroll et al. (10). The validation set was not utilized for the Ridge model, as it does not utilize early stopping, hyperparameter tuning, or best epoch selection.

As shown in Table 1, VIPER surpasses the performance of the previous state-of-the-art model, ProSmith, by 29.6% (McNemar’s test P-value: <0.0001 (45)), achieving a Matthews Correlation Coefficient (MCC) of 0.407 on unseen substrates and similarly outperforming ProSmith on F1 Score by 17%, ROC AUC by 4.1%, PR AUC by 51.6%, and Accuracy by 2.5%.

**Table 1:**
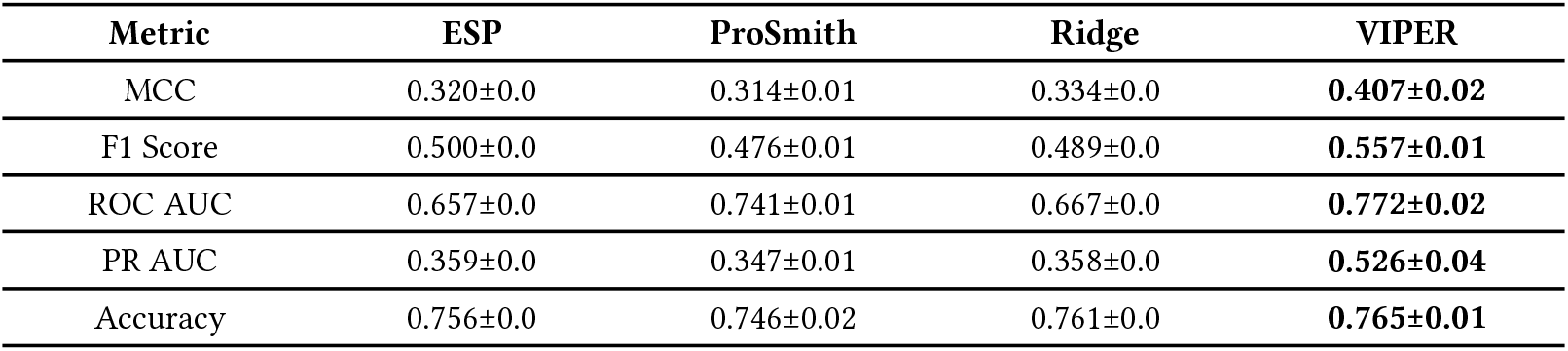
Our model compared with previously introduced models on unseen substrates. The shown score is the mean of three runs. The best score is bolded. All models were run in triplicate to compute the standard error with the exception of ESP, and Ridge, as they are deterministic.

| Metric | ESP | ProSmith | Ridge | VIPER |
| --- | --- | --- | --- | --- |
| MCC | 0.320±0.0 | 0.314±0.01 | 0.334±0.0 | **0.407±0.02** |
| F1 Score | 0.500±0.0 | 0.476±0.01 | 0.489±0.0 | **0.557±0.01** |
| ROC AUC | 0.657±0.0 | 0.741±0.01 | 0.667±0.0 | **0.772±0.02** |
| PR AUC | 0.359±0.0 | 0.347±0.01 | 0.358±0.0 | **0.526±0.04** |
| Accuracy | 0.756±0.0 | 0.746±0.02 | 0.761±0.0 | **0.765±0.01** |

### 2.4 Out-of-distribution benchmarking methodology

To evaluate the out-of-distribution generalization performance of enzyme-substrate prediction models we broke down our dataset into k folds, where k is the number of enzyme families; we then trained the model on k - 1 folds where the fold subtracted was then used as the test set, in a manner similar to a typical KFold cross-validation. For VIPER the performance reported for each fold is the model’s best performance out of ten epochs, the best epoch is selected using a in-distribution validation set consisting of 10% of the training set.

This methodology ensures minimal data similarity between the folds, as almost no substrates are shared between the groups and the catalytic mechanism of the enzymes is inherently different. This is exemplified by Figure 3 where we find that our benchmarking methodology significantly reduces sequence similarity between the train set and test set enzymes with an average 601.312% decrease in BLOSUM62 Mean Similarity of Matched Residues (Mann-Whitney U test: p-value <0.0001) and a 3.753% decrease in Harmonic Mean of Fraction of Matched Residues (Mann-Whitney U Test: p-value <0.0001).

**Figure 3.**
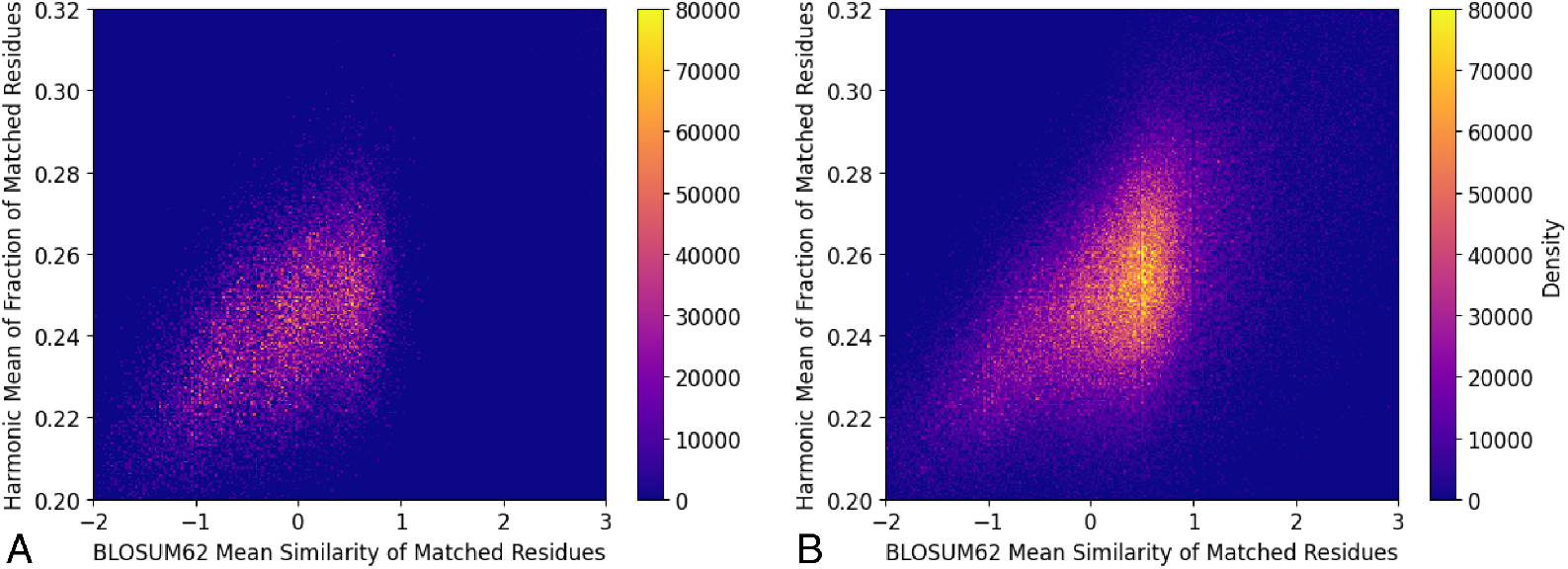
Comparison between our splitting methodology and a typical 80/20 train-test split with 40% sequence similarity cutoff. Every enzyme sequence in the test set was compared to every enzyme sequence in the train set using the Needleman–Wunsch global alignment algorithm (46). **A**. Enzyme family splitting methodology on Esterases (Our splitting methodology) **B**. 80/20 train-test split with 40% sequence similarity cutoff (Previous splitting methodology).

Additionally, we find that our splitting methodology lowers the mean Tanimoto coefficient by 45.3% (Supplementary Table 2) showing that our splitting methodology significantly reduces both enzyme similarity and molecule similarity. To further ensure that there is minimal data similarity we used a K-Nearest Neighbors (KNN) model to see how well a purely memorization-driven approach performs. Additionally, all models were run in triplicate to be able to calculate standard error.

### 2.5 VIPER exhibits superior generalization performance on out-of-distribution data compared to prior models

As shown in Table 2, VIPER demonstrates superior performance compared to ProSmith (11), ESP (10), KNN, and Ridge (12) in our out-of-distribution benchmark. The baseline K-Nearest Neighbors model, despite its simplicity, outperforms ProSmith, ESP, and Ridge, and achieves the highest performance on the BKACE and aminotransferase sets (albeit with relatively low MCC scores of 0.084 and 0.056 respectively). The observation that previous models underperform a simple nearest-neighbor approach suggests limitations in their ability to learn generalizable enzyme-substrate interactions. While the KNN outperforms Ridge, ESP, and ProSmith it still generally performs poorly in absolute values further validating that our benchmark effectively minimizes data leakage. VIPER achieves an average 141.463% improvement (McNemar’s test P-value: <0.0001 (45)) over KNN. ESP performs near random chance on most folds, while the Ridge Regression and ProSmith models, while they perform better than ESP, still show poor generalization across enzyme families. It is noteworthy that all models, including VIPER, show particularly poor performance on the phosphatase enzyme family. This can be attributed to phosphatases comprising 44.4% of our training dataset; when this substantial portion is removed for evaluation, model performance deteriorates significantly as detailed in Supplementary Table 1.

**Table 2:** Enzyme-substrate prediction models were evaluated on an unseen enzyme family benchmark. Each model was run in triplicate, and the mean score, along with its standard error, is reported. The best MCC score is bolded. Supplementary tables 4 through 6 provide additional results, including F1 Score, ROC AUC, and PR AUC, in addition to the MCC score reported here.

| Dataset | KNN | ProSmith | ESP | Ridge | VIPER |
| --- | --- | --- | --- | --- | --- |
| Phosphatases | 0.015±0.0 | 0.02±0.01 | 0.01±0.0 | 0.094±0.0 | -0.01±0.01 |
| Esterases | 0.084±0.0 | 0.03±0.0 | 0.001±0.0 | 0.023±0.0 | **0.110±0.03** |
| GT Acceptors | 0.07±0.0 | 0.011±0.04 | -0.029±0.0 | 0.031±0.0 | **0.160±0.04** |
| GT Donors | -0.058±0.0 | 0.0±0.0 | -0.057±0.0 | 0.0±0.0 | **0.120±0.025** |
| BKACE | **0.084±0.0** | 0.029±0.0 | -0.026±0.0 | 0.058±0.0 | 0.0±0.01 |
| Thiolasases | 0.161±0.0 | 0.024±0.0 | -0.067±0.0 | -0.04±0.0 | **0.256±0.03** |
| Halogenases | -0.023±0.0 | 0.026±0.0 | -0.004±0.0 | 0.058±0.0 | **0.190±0.02** |
| Aminotransferases | **0.056±0.0** | -0.003±0.02 | 0.0±0.0 | 0.194±0.0 | 0.036±0.02 |
| Nitrilases | -0.02±0.0 | 0.011±0.0 | -0.005±0.0 | -0.121±0.0 | **0.032±0.01** |
| Mean | 0.041±0.06 | 0.016±0.01 | -0.017±0.025 | 0.033±0.08 | **0.099±0.09** |

### 2.6 VIPER can express confidence

Discriminating between results with high and low confidence is crucial for a model intended for practical use. While there are various techniques to assess the uncertainty of a deep learning model such as Monte Carlo Dropout (47), Deep Ensembles (48), and Bayesian Neural Networks (49), we opted to use Monte Carlo dropout due to it’s simplicity and computational efficiency. Monte Carlo dropout is a technique in which the model’s dropout layers are enabled during inference; this causes the model to produce slightly different predictions for every forward pass, creating a distribution of predictions. Once a distribution is created we derive it’s confidence using the inverse entropy of the distribution (Supplementary Equation 1).

We evaluated the efficacy of this technique in two ways. First, we created two evaluation datasets with differing levels of difficulty to assess how well the confidence scores reflect the model’s predictive uncertainty. The normal dataset employed a standard 80/10/10 train, validation, test random split. In contrast, the difficult dataset was created by holding out the esterase enzyme family, presenting a more challenging scenario where the model must generalize to enzymes with different catalytic mechanisms. Secondly, we analyzed the relationship between VIPER’s confidence and performance by plotting cumulative confidence thresholds against various performance metrics (Figure 4A); this demonstrates how the model’s confidence scores correlate with its predictive performance, providing insight into whether the confidence scores can be used as reliable indicators of prediction quality.

**Figure 4.**
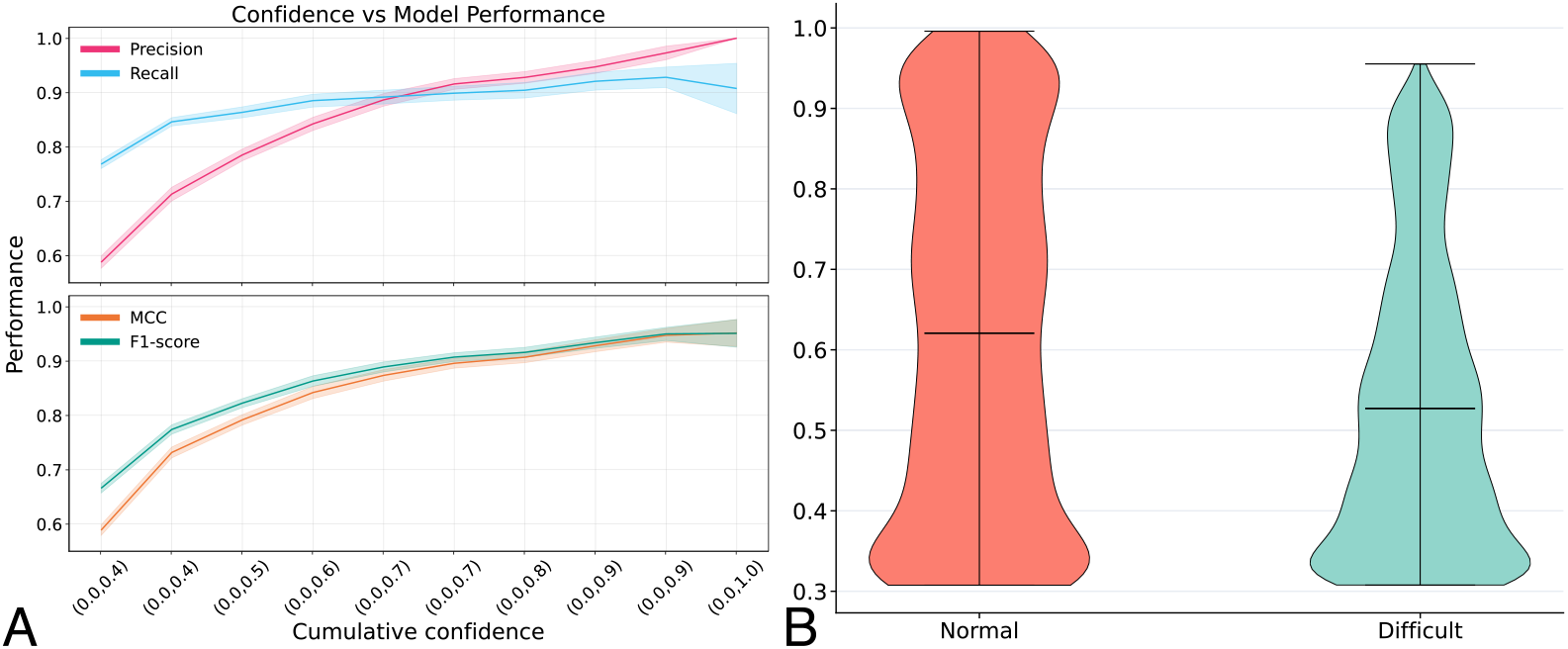
Analyzing the efficacy of using Monte Carlo dropout derived inverse entropy as a confidence metric for VIPER. **A**. Analysis of VIPER’s performance as a function of cumulative confidence thresholds. Shown are the MCC, F1-Score, Precision, and Recall metrics. **B**. Comparison of VIPER confidence score distributions between normal test conditions (standard 80/10/10 random split) and difficult test conditions (held-out esterase enzyme family).

As evident from Figure 4A, as VIPER’s confidence increases, so do the MCC, F1-score, precision, and recall. Each of these metrics increases by 58%, 39%, 11%, and 69%, respectively, between a cumulative confidence of 0.4 and 1.0, and in Figure 4B, we observe that the confidence scores of the difficult test set are significantly lower than those of the normal set. The mean confidence of the normal set is 0.621, while the mean confidence of the difficult set is 0.527, a 15.1% decrease in the mean confidence of the difficult set compared to the normal set.

### 2.7 Component-wise ablation study reveals role of cross-attention

To evaluate the contribution of each architectural component to VIPER’s performance, we conducted a comprehensive ablation study by removing components one at a time while keeping the rest of the architecture intact. We focused on four critical components: the cross-attention layer (−CROSS) which enables protein-molecule interactions, the LSTM layer (−LSTM) which captures sequential dependencies, the convolutional neural networks (−CNNs) which extract features from the input embeddings, and the SAM optimizer (−SAM) which promotes better generalization through flat minima optimization. We evaluated each ablated version against both the unseen molecule benchmark, and the out-of-distribution enzyme family benchmark to understand how each component contributes to VIPER’s performance and generalization capabilities.

As shown in Table 3, we observe a significant performance regression upon removing any of the model’s components. The largest performance regression occurs when removing the CNNs showing that they are important for extracting features to be used by the cross-attention layer. The second largest performance regression occurs upon removal of the cross-attention layer; significantly harming the model’s ability to learn CPIs; removing it leads to a decrease in the MCC by 24.3%, the F1 Score by 12.2%, the ROC AUC by 6.6%, the PR AUC by 7.4%, and the accuracy by 3.1%. These findings further support the notion that VIPER is not merely acting as an advanced nearest-neighbor model, but rather learning broad enzyme-substrate interactions. Moreover, this study demonstrates an improvement over the results obtained by Goldman et al. (12), where models without CPI mechanisms either outperformed or were equivalent to those with CPI mechanisms.

**Table 3:**
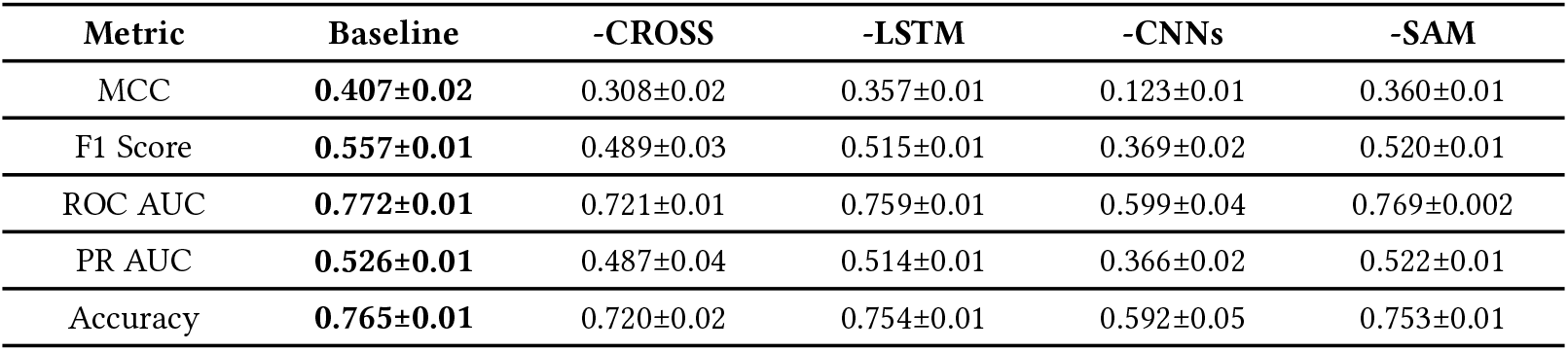
Results of ablation study involving the removal of different components of VIPER was conducted, and its performance is compared to the baseline. The mean performance across three runs is presented along with its standard error. Performance is assessed using the distant unseen molecule benchmark. The best score in each row is bolded.

| Metric | Baseline | -CROSS | -LSTM | -CNNs | -SAM |
| --- | --- | --- | --- | --- | --- |
| MCC | **0.407±0.02** | 0.308±0.02 | 0.357±0.01 | 0.123±0.01 | 0.360±0.01 |
| F1 Score | **0.557±0.01** | 0.489±0.03 | 0.515±0.01 | 0.369±0.02 | 0.520±0.01 |
| ROC AUC | **0.772±0.01** | 0.721±0.01 | 0.759±0.01 | 0.599±0.04 | 0.769±0.002 |
| PR AUC | **0.526±0.01** | 0.487±0.04 | 0.514±0.01 | 0.366±0.02 | 0.522±0.01 |
| Accuracy | **0.765±0.01** | 0.720±0.02 | 0.754±0.01 | 0.592±0.05 | 0.753±0.01 |

As can be seen in Table 4, we observe comparable outcomes in the OOD generalization benchmark as for the distant molecule benchmark. While ablated models occasionally outperform the baseline model on specific sets, they underperform the baseline model across all sets.

**Table 4:**
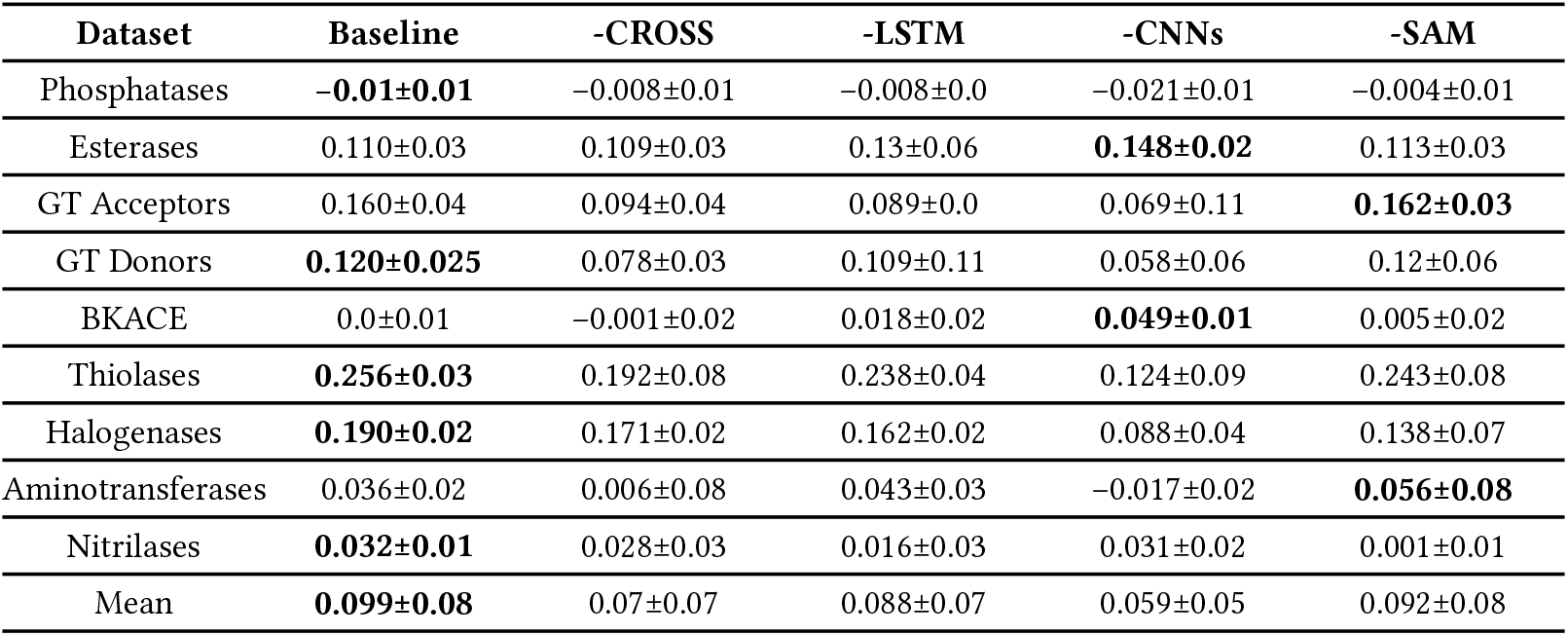
An ablation study was conducted to remove various components of VIPER and compare its performance to the baseline using the unseen enzyme family benchmark. Each model was run in triplicate, and the mean score along with its standard error is reported. The best MCC score is bolded. Supplementary tables 7 through 9 provide additional results, including F1 Score, ROC AUC, and PR AUC, in addition to the MCC score reported here.

| Dataset | Baseline | -CROSS | -LSTM | -CNNs | -SAM |
| --- | --- | --- | --- | --- | --- |
| Phosphatases | **-0.01±0.01** | -0.008±0.01 | -0.008±0.0 | -0.021±0.01 | -0.004±0.01 |
| Esterases | 0.110±0.03 | 0.109±0.03 | 0.13±0.06 | **0.148±0.02** | 0.113±0.03 |
| GT Acceptors | 0.160±0.04 | 0.094±0.04 | 0.089±0.0 | 0.069±0.11 | **0.162±0.03** |
| GT Donors | **0.120±0.025** | 0.078±0.03 | 0.109±0.11 | 0.058±0.06 | 0.12±0.06 |
| BKACE | 0.0±0.01 | -0.001±0.02 | 0.018±0.02 | **0.049±0.01** | 0.005±0.02 |
| Thiolases | **0.256±0.03** | 0.192±0.08 | 0.238±0.04 | 0.124±0.09 | 0.243±0.08 |
| Halogenases | **0.190±0.02** | 0.171±0.02 | 0.162±0.02 | 0.088±0.04 | 0.138±0.07 |
| Aminotransferases | 0.036±0.02 | 0.006±0.08 | 0.043±0.03 | -0.017±0.02 | **0.056±0.08** |
| Nitrilases | **0.032±0.01** | 0.028±0.03 | 0.016±0.03 | 0.031±0.02 | 0.001±0.01 |
| Mean | **0.099±0.08** | 0.07±0.07 | 0.088±0.07 | 0.059±0.05 | 0.092±0.08 |

### 2.8 A web server for easy use of VIPER

We implemented a web server for easy use of VIPER. It is available at https://viperwebserver.com/; the web server takes a SMILES string (50) and a protein amino acid sequence as input, and produces a prediction as well as a confidence value. Additionally, the server allows users to upload a CSV with up to 100 rows for bulk predictions. We recommend utilizing our open-source scripts for large bulk runs.

## 3 Discussion

Here we introduce VIPER, a novel machine learning model for enzyme-substrate prediction that outperforms the previously proposed models: ESP (10), ProSmith (11), and Ridge Regression (12). Our model achieves an average MCC of 0.407 on our unseen substrate benchmark, representing a 29.6% improvement over the previous state-of-the-art model, ProSmith. This substantial performance improvement represents a significant step toward practical in-silico prediction of enzyme substrates.

We have also introduced a novel benchmarking methodology to better understand the out-of-distribution generalization capabilities of enzyme-substrate prediction models. We achieved this by stratifying enzymes in our dataset into families, which is significantly more effective at minimizing enzyme-substrate pair similarity than sequence similarity, due to the inherent differences in catalytic function between enzyme families. Additionally this method significantly reduces molecule similarity where we observed a 45.3% drop in the mean Tanimoto coefficient.

Additionally we have shown that previous datasets introduced for enzyme-substrate prediction have significant flaws that make them largely unsuitable for the training of enzyme-substrate prediction models capable of generalizing to unseen molecules and enzymes; we introduced a novel high-quality dataset for enzyme-substrate prediction, building upon the work for Goldman et al. (12), we performed extensive filtering, reconstruction, and standardization of the Goldman dataset to produce a high quality dataset of enzyme-substrate reactions. While this dataset lacks diversity it’s annotations are of significantly higher quality.

Despite the significant performance improvement VIPER provides over previous models, the model’s performance is still suboptimal for practical use, and like all current enzyme-substrate prediction models should not be used for enzyme families not included in the training data. To further improve model performance, future work could explore the use of semi-supervised learning to increase the available training data or the extraction of data from databases such as BRENDA (38), Sabio-RK (51), or by extracting data directly from publications. Acquiring more diverse data is likely to lead to a large improvement in performance as our current dataset contains only a narrow selection of enzymatic reactions and enzyme families, one key area where our dataset is deficient is coupling reactions performed by enzymes like ligases, synthetases, and polymerases. VIPER is unlikely to generalize well to these enzymes as the reactions they catalyze are distant from anything in our training data. The downside to acquiring data from Sabio-RK or BRENDA is it will have to be cleaned extensively as this data is of quite low quality due to misannotations, high sparsity, human error in curation, and high experimental variance (39–41). Another possible approach for improving model performance is to integrate physical principles of molecule-protein interaction into enzyme-substrate prediction models, which could lead to further improvements in model performance (52,53). The addition of 3D protein information could also be explored as this is critical for determining substrate preferences in many enzyme-catalyzed reactions (54–57).

The successful development of a model that can generalize to new enzyme families and substrates has great potential for industrial synthesis. By utilizing the ability of enzymes to be highly regiospecific, stereospecific, and chemospecific complex synthetic processes can be greatly simplified (1,2), resulting in higher yields and fewer synthesis steps compared to traditional chemical synthesis. This is exemplified by the biosynthesis of QS-21, an FDA-approved immunostimulatory adjuvant commonly used in vaccines (4). The biosynthesis of QS-21 requires 20 steps (3) in contrast to the chemical synthesis, which requires 76 steps (5–7), significantly improving the yield and potentially decreasing the manufacturing cost. Enzymes have also been used to simplify the synthesis of several FDA-approved pharmaceuticals, such as Montelukast (58), Sitagliptin (59,60), Pregabalin (61), and many other pharmaceuticals and high-value compounds (1). Furthermore, VIPER can aid in understanding biological systems by increasing our understanding of substrates for uncharacterized enzymes in various organisms, providing new insights into metabolic pathways and disease mechanisms.

## 4 Methods

### 4.1 Software

Machine learning models were implemented in PyTorch (62) in Python (63). XGBoost (13) was used for gradient-boosting tree models.

### 4.2 Acquisition of data

We acquired the sets: phosphatases (31), esterases (32), glycosyltransferases (20), beto-keto acid cleavage enzymes (33), thiolases (34), halogenases (35), aminotransferases (36), and nitrilases (37) from the pre-processed data made available by Goldman et al. (12).

The selection of enzyme families for inclusion in our dataset was based on several criteria: data accessibility, the presence of non-proprietary enzymes, and overall data quality. This approach ensured a relatively diverse representation of enzyme classes and reaction types, thereby enhancing the broad applicability of our model. The final dataset comprised approximately 51,000 experimental observations. It is worth noting that the data for the phosphatases set was significantly more extensive than data for other sets, constituting 44.4% of the dataset.

### 4.3 Preprocessing of data

We found that the data for the BKACE set (33) made available by Goldman et al. (12) did not include the full enzyme sequence, unlike the other sets but instead just the results of a multi-sequence alignment. To resolve this we acquired the raw sequences with Uniprot IDs from the source paper’s supplementary (33) and the raw MSA data with Uniprot IDs from the unprocessed data made available by Goldman et al. (12). We then used this information to map the correct amino acid sequences onto the BKACE dataset.

Some of the sets used isomeric SMILES strings while others used canonical SMILES strings (50). To resolve this we used RDKit (64) with the Merck molecular force field optimization (65) to generate an initial geometry for the molecule which was then saved to an xyz file. We then used the initial structures as input for xTB (66) with the GFN2 model (67) to perform a semi-empirical geometry optimization in gas-phase with the energy convergence limit set to 5 × 10^−8^hartrees and the gradient convergence set to 5 × 10^−5^ Eh ·α^−1^. We then extracted SMILES strings from these molecular structures using RDKit. Some molecules were excluded as RDKit was unable to determine the appropriate bond order. These SMILES strings were then standardized using the PubChem standardization service via their API (68). We did not process the nitrilase set in the same manner as RDKit was not able to extract an appropriate bond order for a large number of molecules in this set instead the SMILES strings provided by Goldman et al. (12) were canonicalized with RDKit and then standardized with the PubChem standardization service via their API (68). This pre-processing pipeline was applied to all external models in the same manner as it was applied to VIPER.

Activity data for the phosphatases (31), esterases (32), glycosyltransferases (20), beto-keto acid cleavage enzymes (33), thiolases (34), halogenases (35), aminotransferases (36), and nitrilases (37) were binarized according to cutoff values provided by Goldman et al. (12).

### 4.4 Creating enzyme representations

To create the enzyme representations, we used the Ankh base model (450M parameters), a protein language model based on the transformer architecture introduced by Elnaggar et al. (24). Ankh takes the protein amino acid sequence as input and passes it through a modified T5 transformer architecture (69), producing embeddings of shape [Sequence Length, 768] where 768 is the hidden size of the model. We decided to use Ankh because of its state-of-the-art (SOTA) performance and its computational efficiency. Ankh was pre-trained on the UniRef50 database (70) with a masking objective. We used the Ankh implementation provided by Elnaggar et al. on GitHub (24).

### 4.5 Creating molecule representations

To generate molecular representations, we employed Molformer, a chemical language model developed by Ross et al. (25) that utilizes a 12 layer transformer architecture. Molformer takes a SMILES string (50) as input and generates a pooled embedding of shape [768] for each molecule, where 768 is the hidden size of the model. The model was pre-trained on 10% of the ZINC database (26) and 10% of the PubChem (27) database using a masking objective. We used the Molformer implementation provided by Ross et al. on GitHub (25).

### 4.6 Validating external models

We obtained the source code for external models from their public GitHub repositories: (10), (11), and (12). The pretrained ProSmith checkpoint provided by Kroll et al. was used to initialize the weights for all ProSmith evaluations. The performance reported for ProSmith is the best-weighted combination of ESM-1b+ChemBERTa2+cls outputs from the gradient boosting ensemble. The weights for the fine-tuned version of ESM1b that ESP utilizes were downloaded from the Zenodo repo provided by Kroll et al. (10).

Hyperparameter optimization was performed for ESP. For the unseen distant molecule benchmark ESP’s parameters were optimized using the validation set (Supplementary Table 3). For the OOD enzyme family testing hyper-parameter optimization was performed individually for every held-out enzyme family using the training data set. The same hyper-parameter optimization process described by Kroll et al. was used (10). The code for hyperparameter optimization was obtained from the ESP GitHub repository (10).

Since ESP is deterministic, standard error was not calculated for its evaluations. All models were trained using the same dataset as VIPER, and with the same testing methodology. The K-Nearest Neighbors (KNN) model was implemented using Scikit-Learn (71). As input, we used a 1024-dimensional molecular ECFP fingerprint with a radius of three, along with an ESM-1b embedding of the enzyme (15).

### 4.7 Lone cofactor error detection

Our lone cofactor detection algorithm is based on the work of Mukhopadhyay et al. (72), who provide a comprehensive database mapping Enzyme Commission (EC) numbers to potential cofactor scaffolds. For each unique molecule-EC number pair in the dataset, we first determine the presence of valid cofactor scaffolds. If scaffolds are present, we compute the Tanimoto coefficient between each possible scaffold in the EC number tranche and the given molecule. Molecules are represented using 2048-dimensional Extended-Connectivity Fingerprints (ECFP). A molecule-EC number pair is designated as a cofactor if its Tanimoto similarity exceeds 0.8 with any of the scaffolds. This process is applied exclusively to positive entries in the dataset. Examples of molecules excluded as cofactors from the Kroll et al. dataset (10) can be seen in Supplementary Figure 1.

### 4.8 Impossible enzymatic reaction detection

We applied the RetroRules reaction rule database, as provided by Duigou et al. (42), to both the Kroll et al. (10) dataset and the VIPER dataset. For each entry, we extracted all reaction rules corresponding to the given EC number. Then, we evaluated every possible reaction rule for that EC number against the substrate, considering an entry correct if at least one rule was valid. We assessed reaction rule validity using RDKit’s SMARTS reaction rule parser (64). We checked rules in both forward and reverse directions, and a reaction was deemed valid if either direction was applicable to the substrate. This process was exclusively applied to positive dataset samples.

It’s important to note that this method has limitations. The RetroRules database does not include every possible enzymatic reaction; this can lead to both overestimation and underestimation of the error rate. Underestimation can be caused in cases where no reaction rules exist for a given EC number; in these cases, we considered all reactions catalyzed by that EC number to be correct. Overestimation can be caused when some reaction rules exist for the EC number, but they are not comprehensive.

### 4.9 EC number acquisition

For the VIPER dataset, we employed CLEANpred (21) to predict the Enzyme Commission (EC) number for each unique enzyme. In the case of the Kroll et al. (10) dataset, EC numbers were primarily extracted from the corresponding UniProt entries for each protein. However, 15.4% of proteins in the Kroll et al. dataset lacked a listed EC number. For these entries, we utilized CLEANpred (21) to predict the EC numbers.

## 5 Data & Code Availability

All data and code is available at the GitHub repo: https://github.com/maxall41/VIPER-Code. Model weights can be found on Zenodo: https://zenodo.org/records/12151573.

## 6 Supplementary

### 6.1 Supplementary Figures

#### 6.1.1 Supplementary Figure 1

**Figure 5.**
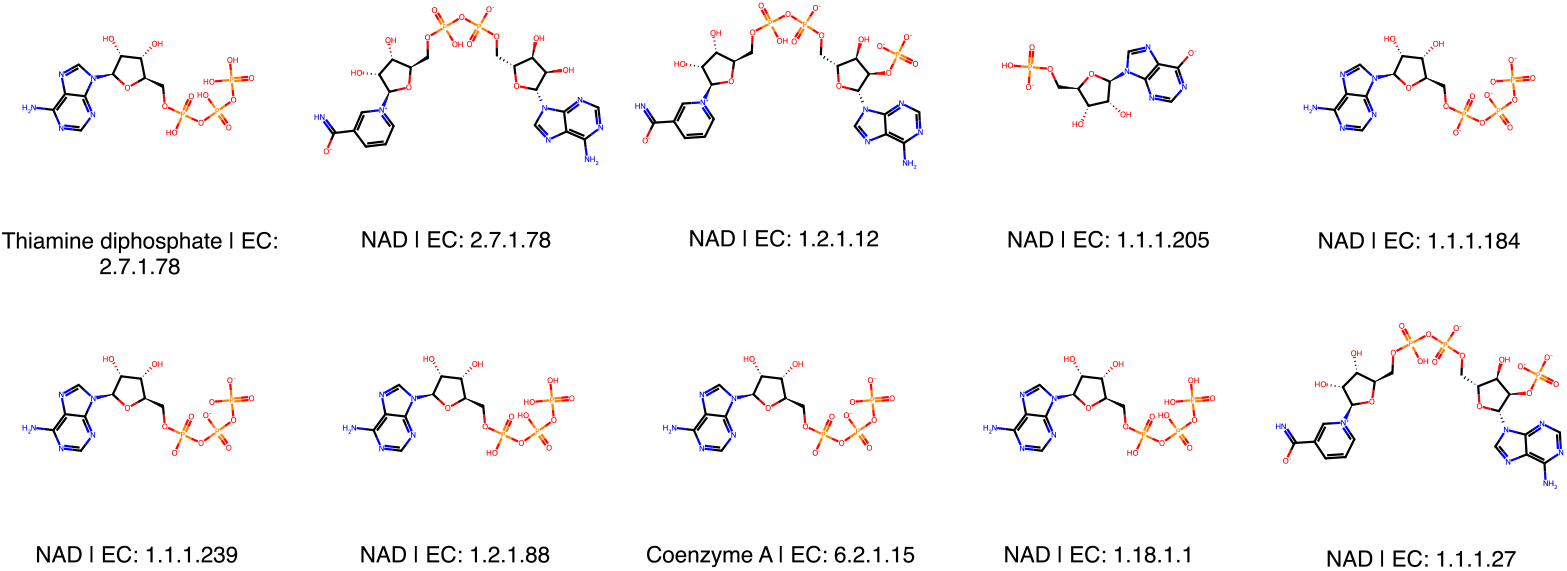
Molecules excluded as cofactors from the Kroll et al. dataset (10).

#### 6.1.2 Supplementary Figure 2

**Figure 6.**
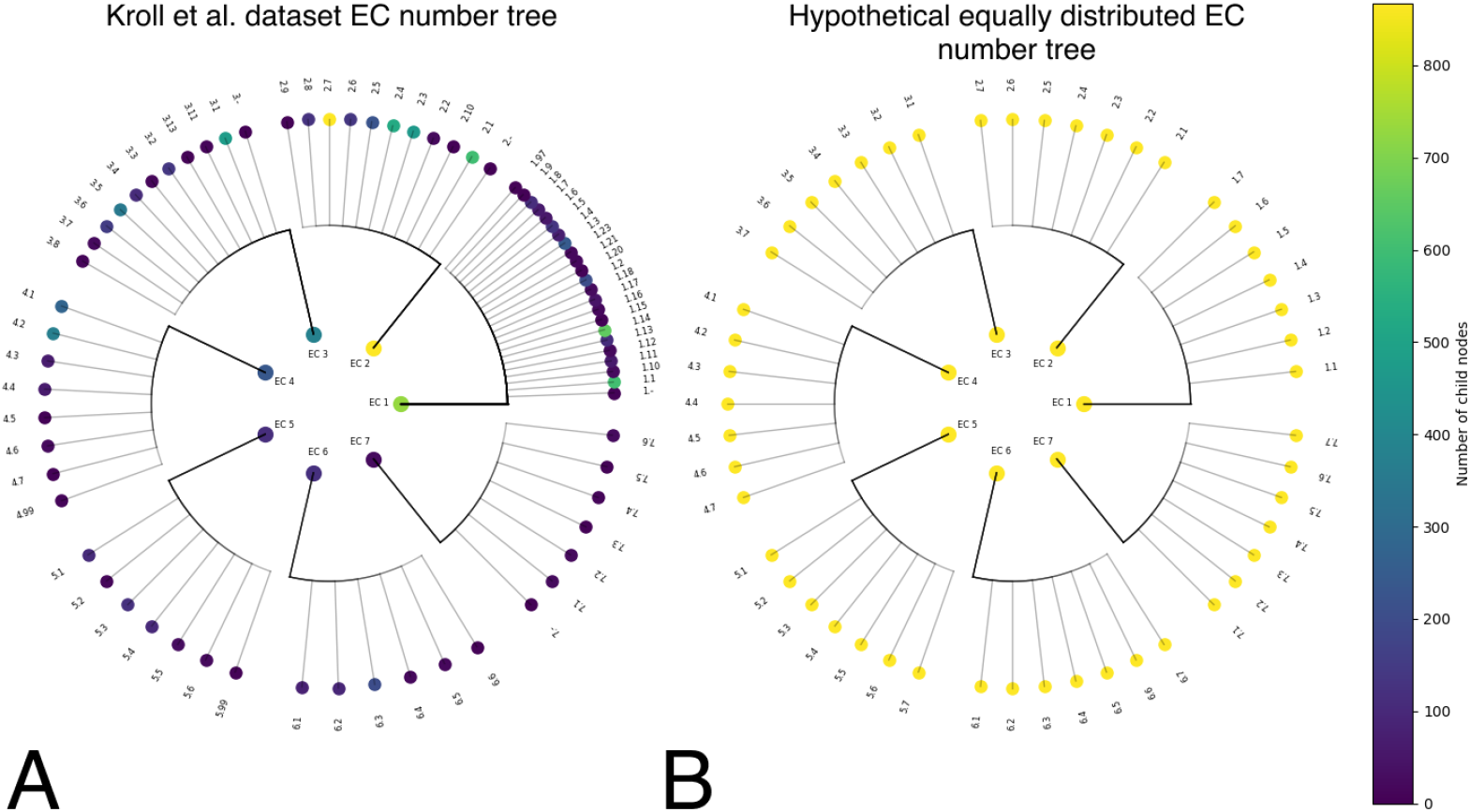
Circular tree of EC number families with the first digit (central nodes) and second digit (outer nodes) of the EC number shown. Color represents the number of proteins under each node. **A**. Distribution of EC numbers in Kroll et al. dataset (10). **B**. Hypothetical even distribution of EC numbers.

### 6.2 Supplementary Tables

#### 6.2.1 Supplementary Table 1

**Table 5:** VIPER Performance with and without phosphatase set compared on unseen enzyme families. Best MCC score is bolded.

| Dataset | VIPER w/phosphatase | VIPER wo/phosphatase |
| --- | --- | --- |
| Esterases | **0.110** | 0.099 |
| GT Acceptors | **0.160** | 0.108 |
| GT Donors | **0.120** | 0.0 |
| BKACE | 0.0 | **0.049** |
| Thiolases | **0.256** | 0.198 |
| Halogenases | **0.190** | 0.0 |
| Aminotransferases | **0.036** | 0.0 |
| Nitrilases | **0.032** | -0.012 |

#### 6.2.2 Supplementary Table 2

**Table 6:** Comparison between mean Tanimoto coefficient for basic 80/20 train/test split (Previous methodology) and enzyme family split (Our methodology). The lowest score is bolded.

| Dataset | Basic Split | Enzyme Family Split |
| --- | --- | --- |
| Phosphatases | 0.138 | **0.089** |
| Esterases | 0.138 | **0.101** |
| GT Acceptors | 0.138 | **0.084** |
| GT Donors | 0.138 | **0.121** |
| BKACE | 0.138 | **0.110** |
| Thiolases | 0.138 | **0.105** |
| Halogenases | 0.138 | **0.073** |
| Aminotransferases | 0.138 | **0.113** |
| Nitrilases | 0.138 | **0.048** |

#### 6.2.3 Supplementary Table 3

**Table 7:** ESP Hyperparameters selected using the unseen molecule benchmark validation set. ESP Hyperparameters were selected using implementation made available in the ESP GitHub repo.

| Name | Value |
| --- | --- |
| learning_rate | 0.05025527658054195 |
| max_delta_step | 3.328719228662338 |
| max_depth | 12 |
| min_child_weight | 2.125754969886389 |
| num_rounds | 375.94371587069827 |
| reg_alpha | 1.7169499373351478 |
| reg_lambda | 1.395084752719229 |
| weight | 0.26980500317480516 |

#### 6.2.4 Supplementary Table 4

**Table 8:** Performance of our model and previously introduced models across different unseen enzyme subsets. The shown score is the mean of three runs. All models were run in triplicate to compute the standard error with the exception of ESP, and Ridge, as they are deterministic. Best F1 score is bolded.

| Dataset | KNN | ProSmith | ESP | Ridge | VIPER |
| --- | --- | --- | --- | --- | --- |
| Phosphatases | 0.151±0.0 | 0.147±0.08 | 0.005±0.0 | **0.304±0.0** | 0.117±0.01 |
| Esterases | 0.159±0.0 | 0.146±0.01 | 0.01±0.0 | 0.203±0.0 | **0.306±0.06** |
| GT Acceptors | 0.323±0.0 | 0.116±0.08 | 0.051±0.0 | 0.017±0.0 | **0.333±0.0** |
| GT Donors | 0.0±0.0 | 0.0±0.0 | 0.0±0.0 | 0.0±0.0 | **0.12±0.03** |
| BKACE | 0.194±0.0 | 0.109±0.06 | 0.0±0.0 | **0.195±0.0** | 0.176±0.0 |
| Thiolases | 0.262±0.0 | 0.027±0.02 | 0.012±0.0 | 0.0±0.0 | **0.513±0.03** |
| Halogenases | 0.04±0.0 | 0.054±0.05 | 0.013±0.0 | 0.155±0.0 | **0.25±0.02** |
| Aminotransferases | 0.011±0.0 | 0.004±0.0 | 0.0±0.0 | **0.36±0.0** | 0.056±0.03 |
| Nitrilases | 0.099±0.0 | 0.014±0.01 | 0.021±0.0 | 0.085±0.0 | **0.104±0.01** |

#### 6.2.5 Supplementary Table 5

**Table 9:** Performance of our model and previously introduced models across different unseen enzyme subsets. The shown score is the mean of three runs. All models were run in triplicate to compute the standard error with the exception of ESP, and Ridge, as they are deterministic. Best ROC AUC score is bolded.

| Dataset | KNN | ProSmith | ESP | Ridge | VIPER |
| --- | --- | --- | --- | --- | --- |
| Phosphatases | 0.507±0.0 | 0.517±0.02 | 0.540±0.0 | **0.575±0.0** | 0.468±0.02 |
| Esterases | 0.525±0.0 | 0.547±0.02 | 0.511±0.0 | 0.578±0.0 | **0.639±0.01** |
| GT Acceptors | 0.543±0.0 | 0.525±0.05 | 0.596±0.0 | 0.619±0.0 | **0.624±0.01** |
| GT Donors | 0.494±0.0 | 0.519±0.11 | 0.527±0.0 | 0.624±0.0 | **0.679±0.03** |
| BKACE | 0.548±0.0 | 0.576±0.03 | 0.500±0.0 | **0.644±0.0** | 0.515±0.02 |
| Thiolases | 0.55±0.0 | 0.58±0.07 | 0.436±0.0 | 0.348±0.0 | **0.679±0.01** |
| Halogenases | 0.49±0.0 | 0.524±0.02 | 0.613±0.0 | 0.607±0.0 | **0.651±0.01** |
| Aminotransferases | 0.503±0.0 | 0.547±0.04 | 0.357±0.0 | **0.643±0.0** | 0.494±0.01 |
| Nitrilases | **0.49±0.0** | 0.508±0.01 | 0.458±0.0 | 0.411±0.0 | 0.487±0.04 |

#### 6.2.6 Supplementary Table 6

**Table 10:** Performance of our model and previously introduced models across different unseen enzyme subsets. The shown score is the mean of three runs. All models were run in triplicate to compute the standard error with the exception of ESP, and Ridge, as they are deterministic. Best PR AUC score is bolded.

| Dataset | KNN | ProSmith | ESP | Ridge | VIPER |
| --- | --- | --- | --- | --- | --- |
| Phosphatases | 0.171±0.0 | 0.172±0.0 | 0.168±0.0 | **0.19±0.0** | 0.16±0.0 |
| Esterases | 0.238±0.0 | 0.229±0.0 | 0.224±0.0 | 0.228±0.0 | **0.298±0.01** |
| GT Acceptors | 0.22±0.0 | 0.199±0.01 | 0.207±0.0 | 0.206±0.0 | **0.259±0.0** |
| GT Donors | 0.298±0.0 | 0.298±0.0 | 0.298±0.0 | 0.298±0.0 | **0.434±0.02** |
| BKACE | **0.119±0.0** | 0.109±0.0 | 0.105±0.0 | 0.108±0.0 | 0.111±0.01 |
| Thiolases | 0.543±0.0 | 0.509±0.0 | 0.505±0.0 | 0.507±0.0 | **0.696±0.02** |
| Halogenases | 0.074±0.0 | 0.079±0.0 | 0.075±0.0 | 0.084±0.0 | **0.158±0.01** |
| Aminotransferases | 0.416±0.0 | 0.413±0.0 | 0.413±0.0 | **0.467±0.0** | 0.42±0.01 |
| Nitrilases | 0.122±0.0 | 0.125±0.0 | 0.124±0.0 | 0.115±0.0 | **0.128±0.01** |

#### 6.2.7 Supplementary Table 7

**Table 11:**
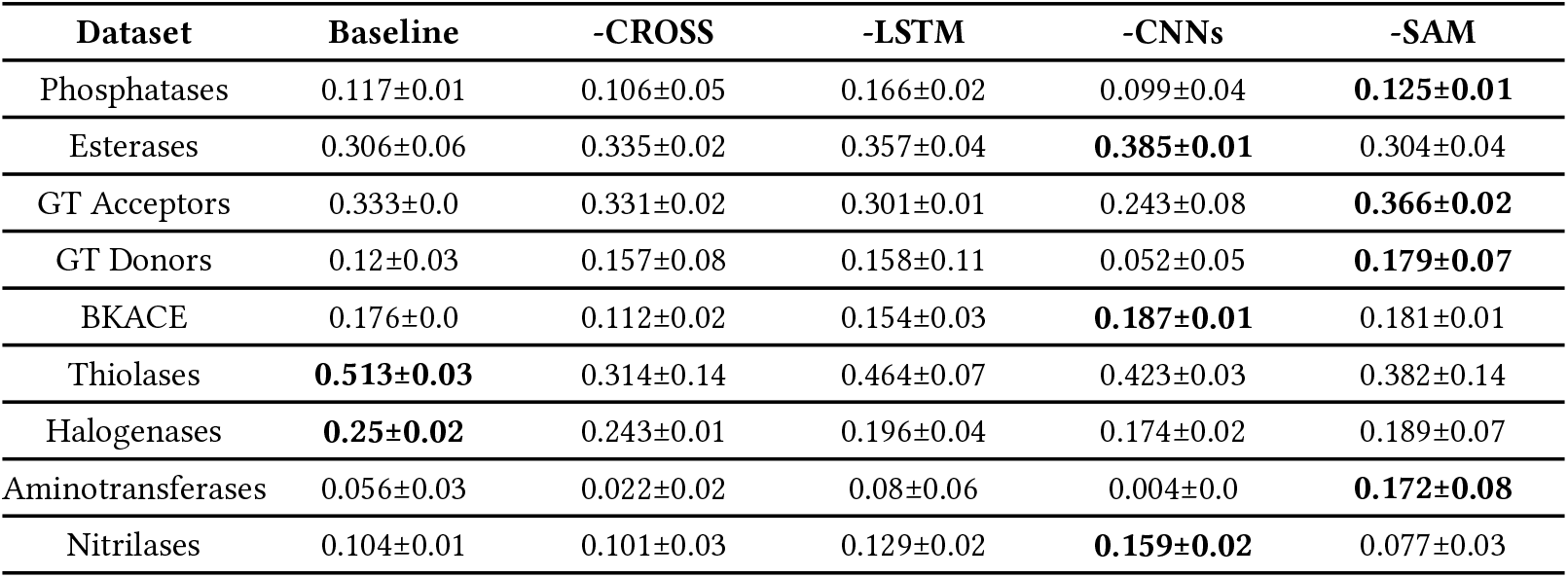
An ablation study was conducted to remove various components of VIPER and compare its performance to the baseline using the unseen enzyme family benchmark. Each model was run in triplicate, and the mean score along with its standard error is reported. The best F1 Score is bolded.

| Dataset | Baseline | -CROSS | -LSTM | -CNNs | -SAM |
| --- | --- | --- | --- | --- | --- |
| Phosphatases | 0.117±0.01 | 0.106±0.05 | 0.166±0.02 | 0.099±0.04 | **0.125±0.01** |
| Esterases | 0.306±0.06 | 0.335±0.02 | 0.357±0.04 | **0.385±0.01** | 0.304±0.04 |
| GT Acceptors | 0.333±0.0 | 0.331±0.02 | 0.301±0.01 | 0.243±0.08 | **0.366±0.02** |
| GT Donors | 0.12±0.03 | 0.157±0.08 | 0.158±0.11 | 0.052±0.05 | **0.179±0.07** |
| BKACE | 0.176±0.0 | 0.112±0.02 | 0.154±0.03 | **0.187±0.01** | 0.181±0.01 |
| Thiolases | **0.513±0.03** | 0.314±0.14 | 0.464±0.07 | 0.423±0.03 | 0.382±0.14 |
| Halogenases | **0.25±0.02** | 0.243±0.01 | 0.196±0.04 | 0.174±0.02 | 0.189±0.07 |
| Aminotransferases | 0.056±0.03 | 0.022±0.02 | 0.08±0.06 | 0.004±0.0 | **0.172±0.08** |
| Nitrilases | 0.104±0.01 | 0.101±0.03 | 0.129±0.02 | **0.159±0.02** | 0.077±0.03 |

#### 6.2.8 Supplementary Table 8

**Table 12:** An ablation study was conducted to remove various components of VIPER and compare its performance to the baseline using the unseen enzyme family benchmark. Each model was run in triplicate, and the mean score along with its standard error is reported. The best ROC AUC is bolded.

| Dataset | Baseline | -CROSS | -LSTM | -CNNs | -SAM |
| --- | --- | --- | --- | --- | --- |
| Phosphatases | 0.468±0.02 | **0.566±0.04** | 0.474±0.01 | 0.457±0.01 | 0.469±0.01 |
| Esterases | **0.639±0.01** | 0.574±0.06 | 0.629±0.02 | 0.627±0.0 | 0.634±0.0 |
| GT Acceptors | 0.624±0.01 | 0.612±0.01 | 0.603±0.02 | 0.61±0.02 | **0.629±0.03** |
| GT Donors | 0.679±0.03 | 0.530±0.05 | **0.729±0.01** | 0.578±0.02 | 0.606±0.05 |
| BKACE | 0.515±0.02 | **0.546±0.05** | 0.503±0.01 | 0.542±0.01 | 0.523±0.03 |
| Thiolases | 0.679±0.01 | 0.573±0.07 | 0.679±0.01 | 0.536±0.02 | 0.593±0.04 |
| Halogenases | **0.651±0.01** | 0.525±0.04 | 0.648±0.01 | 0.614±0.01 | 0.634±0.01 |
| Aminotransferases | 0.494±0.01 | 0.534±0.05 | 0.487±0.01 | **0.558±0.04** | 0.52±0.03 |
| Nitrilases | 0.487±0.04 | **0.626±0.05** | 0.48±0.02 | 0.505±0.03 | 0.477±0.04 |

#### 6.2.9 Supplementary Table 9

**Table 13:** An ablation study was conducted to remove various components of VIPER and compare its performance to the baseline using the unseen enzyme family benchmark. Each model was run in triplicate, and the mean score along with its standard error is reported. The best PR AUC is bolded.

| Dataset | Baseline | -CROSS | -LSTM | -CNNs | -SAM |
| --- | --- | --- | --- | --- | --- |
| Phosphatases | 0.16±0.0 | **0.166±0.01** | 0.162±0.0 | 0.154±0.0 | 0.157±0.0 |
| Esterases | 0.298±0.01 | 0.284±0.0 | **0.305±0.02** | 0.288±0.0 | 0.29±0.0 |
| GT Acceptors | 0.259±0.0 | 0.236±0.0 | 0.244±0.01 | 0.252±0.01 | **0.266±0.02** |
| GT Donors | 0.434±0.02 | 0.415±0.03 | **0.493±0.04** | 0.341±0.02 | 0.36±0.04 |
| BKACE | 0.111±0.01 | 0.111±0.01 | 0.105±0.0 | **0.126±0.01** | 0.12±0.01 |
| Thiolases | 0.696±0.02 | 0.675±0.01 | **0.701±0.01** | 0.615±0.02 | 0.658±0.03 |
| Halogenases | 0.158±0.01 | 0.141±0.0 | **0.167±0.01** | 0.119±0.01 | 0.128±0.02 |
| Aminotransferases | 0.413±0.0 | 0.381±0.01 | 0.401±0.0 | **0.432±0.03** | 0.412±0.03 |
| Nitrilases | 0.128±0.01 | 0.121±0.0 | 0.133±0.01 | **0.145±0.02** | 0.133±0.02 |

### 6.3 Supplementary Text

#### 6.3.1 VIPER Training

No hyperparameter optimization was performed for the VIPER model. The AdamW+SAM optimizer learning rate was set to 0.0005, with a beta1 of 0.9 and beta2 of 0.999. The weight decay was set to 0.01. VIPER was trained for 10 epochs.

#### 6.3.2 Proteomic conservation in Kroll et al. dataset

In Figure 6, we show additional evidence of heavy proteomic conservation in the Kroll et al. (10) dataset by creating an enzyme family tree that shows the distribution of proteins across EC numbers. The node color indicates the number of proteins under that node. We also show a hypothetical tree where the proteins are evenly distributed across all possible EC numbers.

#### 6.3.3 Evaluating final models on imine reductases

We evaluate the production-ready models - ESP (10), ProSmith (11), and VIPER - each utilizing their official weights and respective complete training datasets. This evaluation marks a transition from comparing model architectures on the VIPER dataset to assessing final deployment-ready models. We use imine reductases (73) as our test case, as these enzymes are absent from both the VIPER dataset and the Kroll et al. dataset (10). This choice of test data effectively evaluates each model’s generalization capabilities on an unseen enzyme family. To evaluate these models, we employ a combination of metrics, including MCC, F1 Score, ROC AUC, PR AUC, Accuracy, and the Pearson correlation coefficient between predictions and non-binarized assay readouts. For ProSmith, we generated predictions by submitting imine reductase data to the official web server provided by Kroll et al. (11). For ESP, we utilized the original model weights as published by Kroll et al. For VIPER, we employed our published model weights.

As can be seen in Table 14, all models perform poorly on the imine reductase family. They all have an MCC of 0.0 and an F1 score of 0.0. ESP outperforms ProSmith, while VIPER outperforms both ESP and ProSmith. VIPER outperforms ESP in ROC AUC by 42.7%, PR AUC by 25.2%, Accuracy by 1.9%, and Pearson correlation by 335%. However, with a prediction cutoff of 0.5, all models are worse than random chance, but ROC AUC, PR AUC, and Pearson correlation demonstrate that VIPER maintains a correlation between its output score and the assay readout value.

**Table 14:** Comparing ESP, ProSmith, and VIPER on imine reductase enzymes. MCC, F1 Score, ROC AUC, PR AUC, Accuracy, and Pearson correlation coefficient are shown. The best score in each row is bolded unless all scores are equal.

| Metric | ESP | ProSmith | VIPER |
| --- | --- | --- | --- |
| MCC | 0.0 | 0.0 | 0.0 |
| F1 Score | 0.0 | 0.0 | 0.0 |
| ROC AUC | 0.451 | 0.391 | **0.644** |
| PR AUC | 0.520 | 0.464 | **0.651** |
| Accuracy | 0.465 | 0.465 | **0.474** |
| Pearson | -0.1 | -0.109 | **0.235** |

Imine reductase data, protein sequence identifiers, and molecule structures were extracted from tables in the body of the paper (73). Enzyme-substrate reactions were quantified as the percent of substrate converted to product. Assay readout values were binarized with an absolute value cutoff of 90%, as this is considered to be good activity by Wetzl et al (73).

### 6.4 Supplementary Equations

#### 6.4.1 Model confidence (Supplementary Equation 1)

Let *P*_*i*_ be a prediction. Let *𝔼* be the entropy. Let *n* be the number of forward passes.

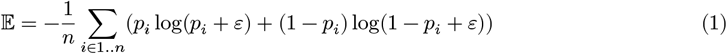

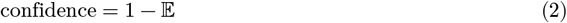

